# Evaluating steroid hormone receptor interactions using the live-cell NanoBRET proximity assay

**DOI:** 10.1101/2023.07.25.550078

**Authors:** Rosemary J Huggins, David Hosfield, Amira Ishag-Osman, Keemin Lee, Elia Ton-That, Geoffrey L. Greene

**Affiliations:** Ben May Department for Cancer Research, University of Chicago, Chicago, IL, United States

**Keywords:** Nuclear receptor, crosstalk, NanoBRET, physical interaction

## Abstract

Steroid hormone receptors play a crucial role in the development and characterization of the majority of breast cancers. These receptors canonically function through homodimerization, but physical interactions between different hormone receptors play a key role in cell functions as well. The estrogen receptor (ERα) and progesterone receptor (PR), for example, are involved in a complex set of interactions known as ERα/PR crosstalk. Here, we developed a valuable panel of nuclear receptor expression plasmids specifically for use in NanoBRET assays to assess nuclear receptor homo- and heterodimerization. We demonstrate the utility of this assay system by assessing ERα/PR physical interaction in the context of the endocrine therapy resistance-associated ERα Y537S mutation. We identify a role of the ERα Y537S mutation beyond that of constitutive activity of the receptor; it also increases ERα/PR crosstalk. In total, the NanoBRET assay provides a novel avenue for investigating hormone receptor crosstalk. Future research may use this system to assess the effects of other clinically significant hormone receptor mutations on hormone receptor crosstalk.

## Background

Steroid hormone receptors are type I nuclear receptors that are implicated in the progression of endocrine-associated cancers, including breast cancer. Approximately 75% of breast cancer cases are characterized as hormone receptor-positive in terms of estrogen receptor (ERα) and/or progesterone receptor (PR)^1^. Androgen receptor (AR), known most for its role in the development of prostate cancer, is also co-expressed in 60-80% of ERα+ breast cancers and is generally an indicator of good prognosis^2–5^. Relatively little is known about the role of mineralocorticoid receptor (MR) in the breast, though recent studies suggest that MR activity may be growth suppressive in breast cancer^6^.

Dimerization is a key step in mediating the function of hormone receptors. Homodimers form more readily than heterodimers due to the high binding affinity between receptors of shared sequence and structure. However, physical interactions between different hormone receptors play an important role in cell function^7–9^. Such physical interactions may occur through a variety of structurally diverse mechanisms that bring different hormone receptors in proximity, including:

1. Heterodimerization, such as the three-point interaction between peroxisome proliferator-activated receptor-ϒ (PPAR-ϒ) and retinoid X receptor (RXR)^10^
2. Allosteric modulation of hormone receptor binding to DNA via DNA binding domain (DBD) interactions^8^
3. Formation of complexes of hormone receptors with shared co-regulators, which are expressed in a temporal and cell-dependent manner^7, 8, 11^

Regardless of the method by which physical interactions between different hormone receptors occur, such interactions play a key role in what is known as hormone receptor crosstalk. Receptor crosstalk can refer to reciprocal gene regulation by two different hormone receptors, hormone-independent activity of a receptor in response to activity by a different receptor, or physical interaction of two receptors in a regulatory complex. For example, ERα/PR crosstalk occurs via:

1. Liganded ERα regulating *PGR* gene transcription^12–16^
2. Liganded PR increasing ERα target gene regulation through ERα phosphorylation^12^
3. PR-dependent chromatin remodeling to facilitate ERα binding^17, 18^
4. ERα/PR physical interaction via regulatory complexes contributing to ligand-independent target gene expression^12, 19, 20^

ERα/PR crosstalk is thought to play a role in breast cancer progression and may contribute to the altered gene expression profile of endocrine therapy (ET)-resistant tumors^12, 17, 19^. ETs such as aromatase inhibitors (AI) or tamoxifen are often the first-line therapy for patients with hormone-sensitive breast cancers and have improved post-surgery outcomes and relapse-free survival^21^. Despite its benefits, ∼25% of patients treated with adjuvant ET for five years or more develop ERα point mutations that drive treatment resistance and contribute to the progression of metastatic breast cancer^22–24^. ERα Y537S is one of the most frequently identified ERα mutations in patients, with the mutation appearing in one-third of circulating tumor cells from blood samples and at least 20% of metastatic tumors^22, 25–28^. Notably, while ERα Y537S is very rarely found in primary treatment-naïve tumors, it is associated with tumor progression, especially in response to aromatase inhibitors, suggesting that ET results in selective pressure toward more resistant and aggressive metastases^26^.

ERα Y537S stabilizes the activating function-2 (AF-2) cleft of the ERα ligand binding domain (LBD) in the agonist-bound conformation, which facilitates constitutive activity of the LBD, even in the absence of ligand binding^29^. Conversely, ERα Y537S alters the antagonist state of AF-2 by reducing the affinity of antagonists for the receptor, thereby increasing resistance to inhibition by selective estrogen receptor modulators and degraders (SERMs and SERDs)^29^. Further investigation into the effects of ERα Y537S on the transcription factor activity of ERα identified ∼900 genes that were significantly induced in ERα Y537S, including several genes that were uniquely bound by ERα Y537S compared to ERα WT^26^.

Given the multimodal nature of ERα/PR crosstalk involving both physical interaction of the receptors through regulatory complexes as well as reciprocal regulation of transcription factor activity, we hypothesized that the functional effects of ERα Y537S are not limited to ERα, but also affect the activity of PR. Here, we utilize the informative NanoBRET assay^30^ for live-cell analysis to determine the effects of ERα Y537S on the physical interaction of ERα and PR. Elucidating the extent to which ERα Y537S alters ERα/PR crosstalk will improve our understanding of how this activating mutation contributes to ET resistance and may offer alternative targets for treating resistant disease. Furthermore, characterization and validation of a full panel of steroid hormone receptor plasmids for NanoBRET assays provide a novel avenue for investigating hormone receptor crosstalk.

## Methods

### Cell lines and growth conditions

HEK293 cells were obtained from the ATCC and maintained in phenol red-free DMEM containing 5% fetal bovine serum (FBS), 1% Pen/Strep, and 1% L-Glutamine. Before NanoBRET assays, HEK293 cells were cultured in phenol red-free DMEM containing 10% charcoal-stripped serum (CSS), 1% Pen/Strep, and 1% L-Glutamine.

### Plasmids and compounds

pCDNA3.1-based plasmids containing the complete coding sequences for the steroid receptor genes were provided by Dr. David Hosfield at the University of Chicago. The compounds NanoBRET Nano-Glo Substrate (Promega #N1571) and HaloTag NanoBRET 618 Ligand (Promega #G9801) were used in NanoBRET assays. Aldosterone (Aldo, Sigma #A9477), 5α- dihydrotestosterone (DHT, Sigma #D-073), progesterone (P4, Sigma #P0130), and β-estradiol (E2, Sigma #E2758) were used in NanoBRET assays. Vehicle (ethanol) was used as a control for all experiments.

### Preparation of NanoLuc and HaloTag plasmids for NanoBRET

N- and C-terminal fusion of the NanoLuc and HaloTag reporters were appended to the steroid receptor genes using Gibson Assembly with primers designed using the assembly tools within the program SnapGene (from Insightful Science; available at snapgene.com). Briefly, PCR was used to amplify the coding regions of the steroid receptor genes and to linearize the expression plasmids pHTN HaloTag CMV Neo or pFLN-1 NanoLuc (Promega #N1811, see Supplemental Table 1 for primer details). PCR products were then isolated via gel electrophoresis and assembled using HiFi assembly mix (NEB #E2621L). Plasmids were verified by DNA sequencing and stored at -20°C until use in NanoBRET assays.

### NanoBRET assay

After culturing HEK293 cells in stripped media (DMEM containing 10% CSS) for 48 hours, cells were trypsinized and collected. Using a Countess cell counter and trypan blue staining at a 1:1 ratio of stain to cell solution, the number of live cells was calculated, and the cell solution was diluted to 1e6 cells/mL in stripped media. Using a multichannel pipette, 100uL of cell solution was dispersed into each well of a 96-well plate (black, clear-bottomed plate) for 1e5 cells/well. After 24 hours, cells were co-transfected with the appropriate HaloTag and NanoLuc plasmids (experimental or control plasmids, at concentrations optimized by preliminary experiments – generally 250ng/uL for HaloTag plasmids and 50ng/uL for NanoLuc plasmids) plus transfection reagent (20uL Lipofectamine 2000 + 800uL PBS) followed by incubation for 24hrs at 37°C and 5% CO2. The following day, cells were treated with the appropriate compounds and 10uL of 500nM HaloTag ligand (G618) for 3 hours. Just before assay quantification in a luminometer, NanoLuc substrate was added to each well, followed by brief shaking to mix. Assays were quantified using the NanoBRET protocol on the TECAN Synergy Neo plate reader in the University of Chicago Cellular Screening Center. This protocol measures total donor luminescence at 450nm (indicative of NanoLuc expression) and total acceptor fluorescence at 610nm (indicative of HaloTag expression). Data is analyzed as the ratio of acceptor fluorescence to donor luminescence (fluorescence/luminescence) as described by Machleidt and colleagues^30^.

### Statistical analysis

Data (except dose-response curves for NanoBRET assays) were analyzed by ordinary two-way ANOVA (α = 0.05) with Tukey’s multiple comparisons tests to compare between treatments within each cell line, as well as between cell lines for each treatment. Dose-response curves were analyzed with nonlinear regression for log(treatment) vs. response to calculate log(IC50) values. Ordinary one-way ANOVA (α = 0.05) with Tukey’s multiple comparisons tests were used to compare IC50 values between each treatment. For all analyses: **** p-value < 0.0001, *** p-value < 0.001, ** p-value < 0.01, * p-value < 0.05.

## Results

### Optimization and validation of nuclear receptor expression plasmids for NanoBRET assays

Prior to utilizing NanoBRET assays to experimentally investigate the effects of various manipulations (ligand treatment, receptor mutations, etc.), the optimal NanoLuc and HaloTag positions were determined through a complete comparison of quantified fluorescence/luminescence ratios for each possible arrangement of C-terminal and N-terminal tag positions (Fig. 1). For each nuclear receptor (ERα, PR-B, AR, and MR), NanoLuc and HaloTag relative positions were considered optimal based on the ability of the nuclear receptors to homodimerize in response to the receptor’s native ligand without interference from the position of the NanoBRET tags (Fig. 2a-d). Briefly, C-terminal HaloTag and NanoLuc positioning was optimal for all nuclear receptors assessed. N-terminal NanoLuc positioning on AR was also non-interfering, thus optimal positioning for AR can be determined according to specific experimental conditions in downstream applications (Fig. 2e).

**Figure 1.**
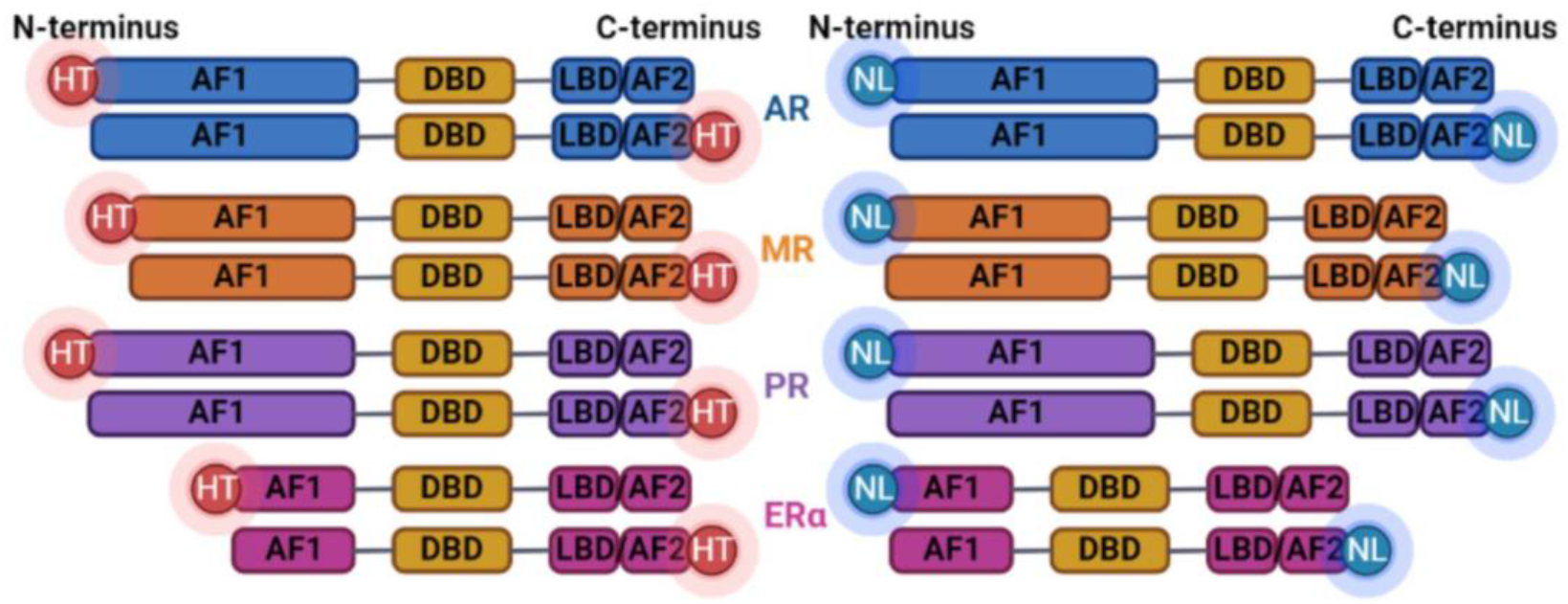
Diagram depicting possible combinations of HaloTag and NanoLuc conformations with each included steroid hormone receptor. From top to bottom: androgen receptor (AR), mineralocorticoid receptor (MR), progesterone receptor (PR), estrogen receptor (ERα). Graphic created with BioRender.com.

**Figure 2.**
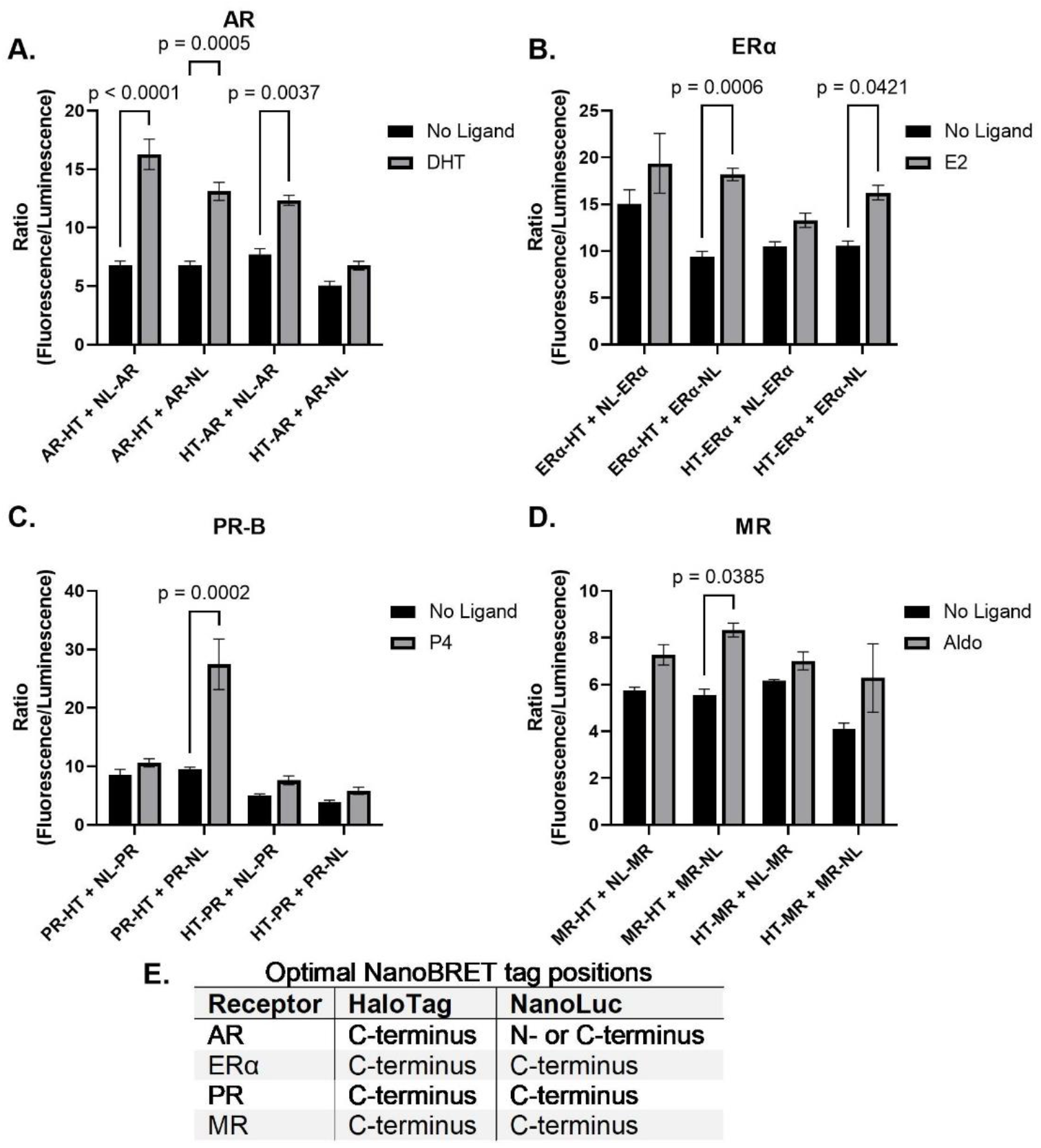
Optimal HaloTag and NanoLuc position allows for ligand-induced homodimerization of nuclear receptors. **A-D.** NanoBRET ratios of fluorescent to luminescent signal quantified upon addition of the NanoLuc substrate to cells treated with vehicle or the native receptor ligand (100nM) for **A)** ERα, **B)** PR-B, **C)** AR, and **D)** MR homodimers. **E)** Table of optimal NanoBRET tag positions based on NanoBRET ratios measured in **A-D**.

To further confirm that receptor homodimerization was not affected by NanoBRET tagging of the receptors, the native ligand of each receptor (as described above) was titrated to assess dose-dependent, ligand-induced nuclear receptor homodimerization. ERα, PR-B, AR, and MR homodimerization in response to E2, P4, DHT, and Aldo (respectively) were strongly dose-dependent, with IC50 values in the nanomolar range (Fig. 3). PR-B, AR, and MR homodimerization was specifically induced in response to their associated native ligands and not others; even at artificially high concentrations of ligand, the nuclear receptors only formed significant proximity-based interactions in response to their own native ligand (Fig. 4). In total, these data highlight the NanoBRET assay as a biologically relevant, live-cell method to quantify proximity-based interactions among hormone receptors.

**Figure 3.**
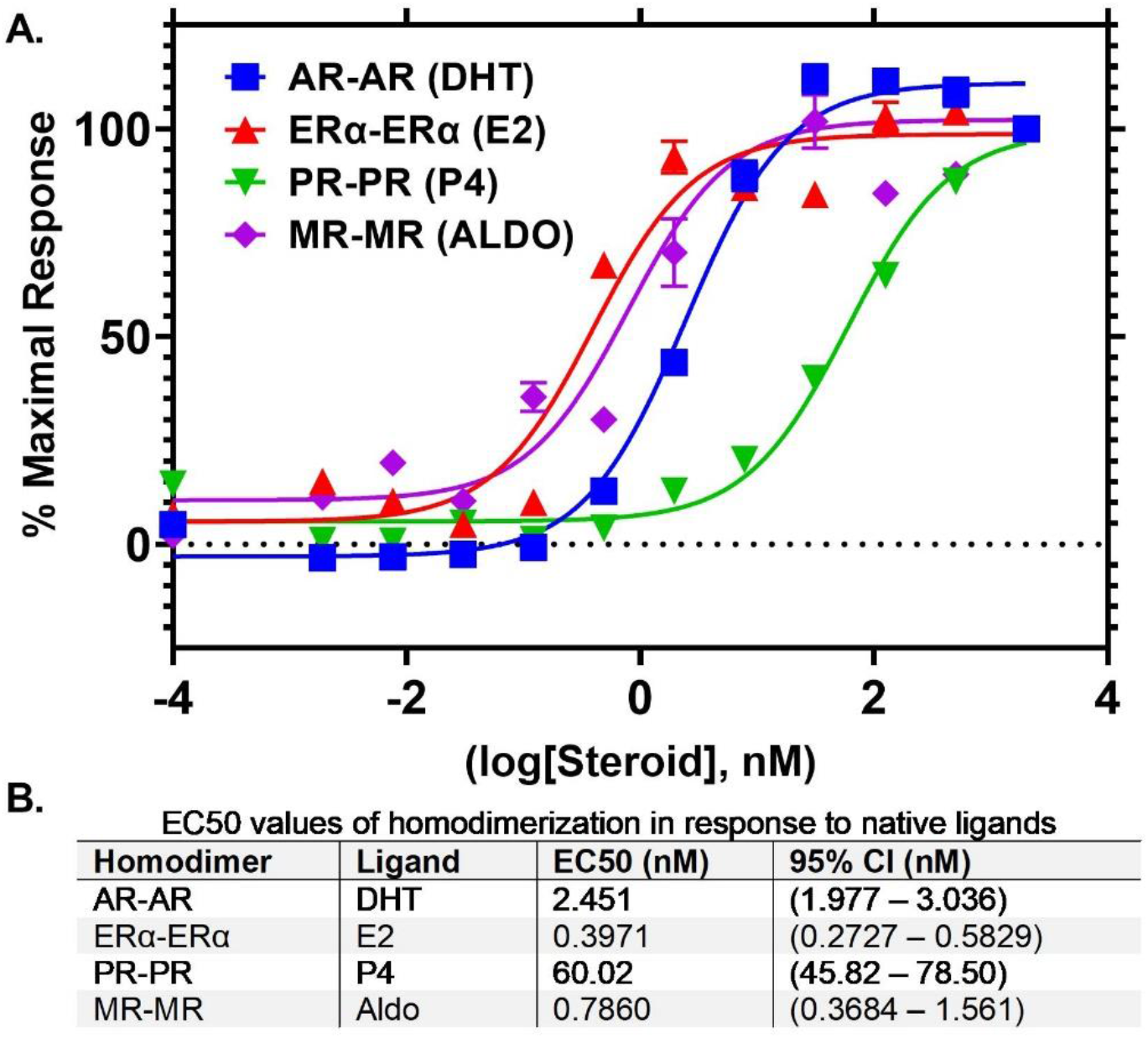
NanoBRET-based nuclear receptor homodimerization is ligand dose-dependent. **A)** Dose-response curves of each NR-HT/NR-NL homodimer pair in response to treatment with their native ligands. **B)** EC50 values calculated from curves in **A**. Data represents minimum 3 biological replicates.

**Figure 4.**
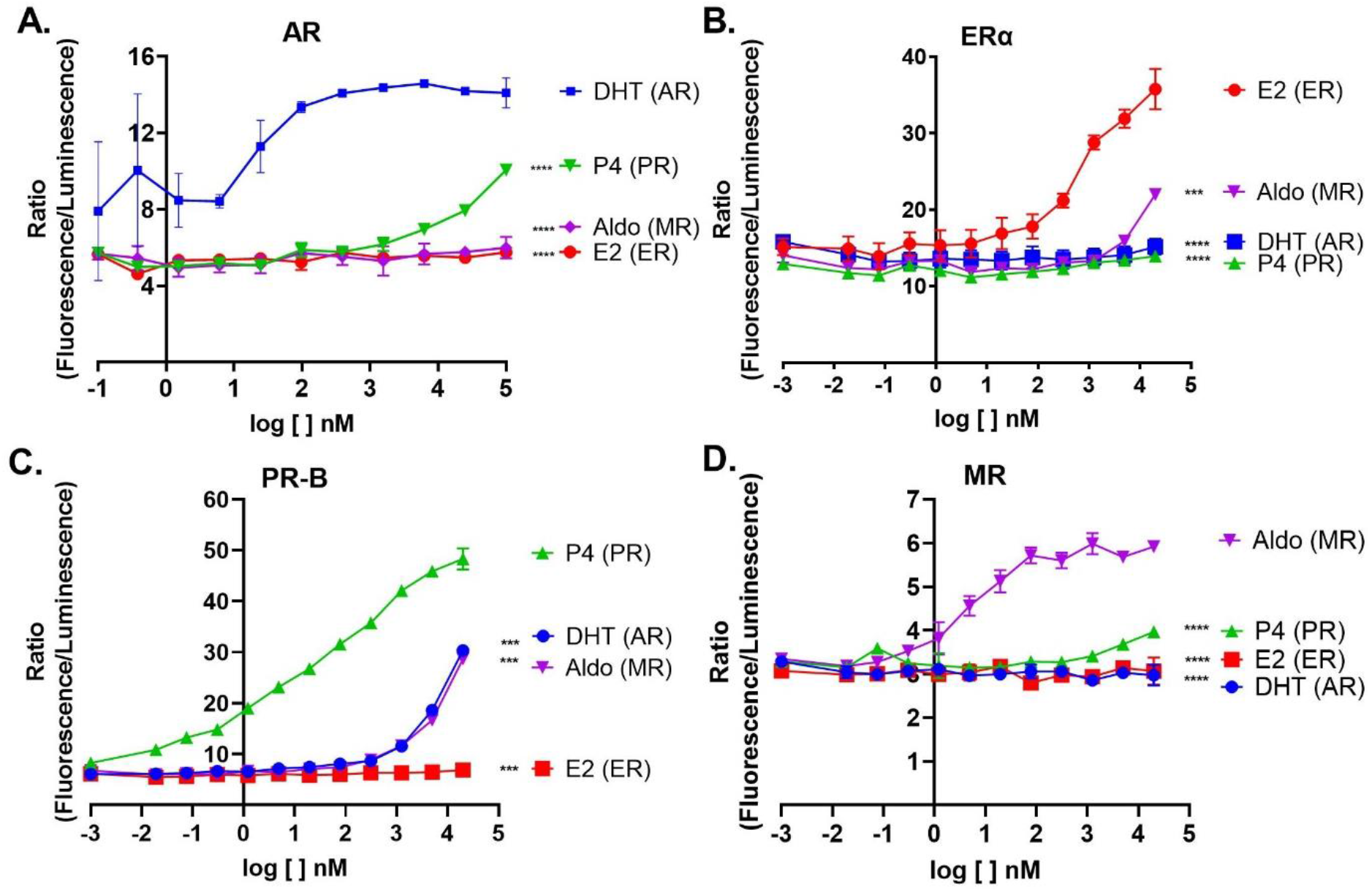
NanoBRET-based nuclear receptor homodimerization is ligand-specific. NanoBRET homodimer formation of **A)** AR, **B)** ERα, **C)** PR-B, and **D)** MR homodimer in response to non-native hormones, relative to formation in response to native ligand. A significant difference between NanoBRET dose-response curves is indicated as *** p < 0.001 or **** p < 0.0001. Data represents minimum 3 biological replicates.

### NanoBRET supports evidence of nuclear receptor heterodimerization, including ERα/PR crosstalk

Upon optimization of the NanoBRET assay for quantifying hormone receptor homodimerization, the method was applied to investigate the proximity-based interaction of ERα with PR-B. As noted previously, physical interaction of ERα and PR-B is a key component of ERα/PR crosstalk^12, 19^. Similar to the optimization of HaloTag and NanoLuc configurations for homodimer formation of each nuclear receptor (Fig. 1, Fig. 2), a methodical approach was taken to determine the optimal configuration of NanoBRET tag positions for assessing proximity-based interactions of ERα and PR-B (Fig. 5a). As with homodimer formation, C-terminal configuration of the NanoBRET tags was optimal, with ERα-HaloTag and PR-B-NanoLuc proximity increasing significantly in response to P4 treatment (Fig. 5b). Optimal tag position and native ligand sensitivity for proximity-based interaction between each nuclear receptor pairing (AR-ERα, AR-MR, AR-PR, ERα-MR, and PR- MR) was also assessed, with several NanoBRET tag configurations proving adequate for each pair (Supp. Fig. 1). Experimental parameters should be considered when determining which tag configuration to use in subsequent experiments.

**Figure 5.**
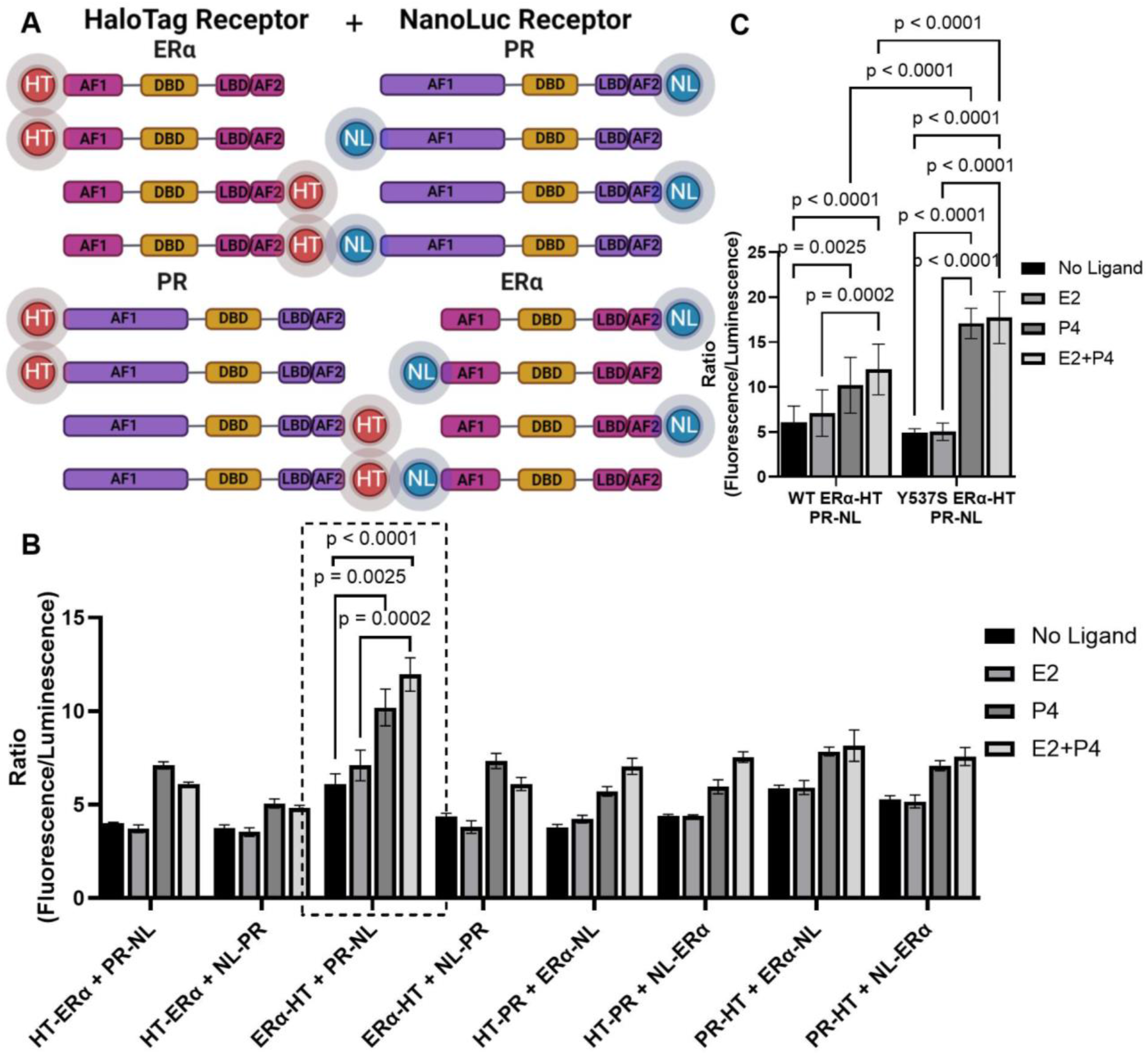
Optimized HaloTag and NanoLuc positioning allows for analysis of ERα and PR proximity-based interaction via NanoBRET assays. **A)** Diagram depicting possible combinations of HaloTag and NanoLuc conformations with ERα and PR. Graphic created with BioRender.com. **B)** NanoBRET ratios of fluorescence to luminescence for each combination of HaloTag and NanoLuc conformations depicted in **A**, in response to vehicle, E2 (ERα native ligand) and P4 (PR native ligand), alone or in combination. Optimal tag positioning based on responsiveness of the receptor proximity to ligand treatment is outlined with a dashed line. **C)** Using optimal NanoLuc/HaloTag positioning, graph shows NanoBRET ratios of ERα WT or ERα Y537S in proximity with PR-B in response to vehicle, E2 (ERα native ligand), and P4 (PR native ligand), alone or in combination. Data represents minimum 3 biological replicates.

### ERα/PR proximity increases in the context of the ERα Y537S mutation

The ERα Y537S mutation is often found in treatment-resistant metastatic breast cancers, and thus it is of significant interest to fully characterize the phenotypic effects of the mutation as well as how it may be targeted. Given the reported role of ERα/PR crosstalk in breast cancer progression and the apparent value of the NanoBRET method for assessing ERα and PR-B proximity-based interactions, we introduced the specific TAT>TCT point mutation in exon 8 of the *ESR1* plasmid to create the ERα Y537S tyrosine to serine amino acid substitution. ERα proximity to PR-B increased significantly in response to progesterone, and this increase was nearly two-fold greater in the context of the ERα Y537S mutation (Fig. 5c). These findings suggest a PR-driven physical interaction between ERα and PR that is enhanced in the context of the treatment resistance-associated ERα Y537S mutation.

## Discussion

Although nuclear receptors canonically function through homodimerization, recent research has suggested that receptor crosstalk may amplify or dampen the activities of nuclear receptors, including those highly implicated in breast cancer^7–12, 18, 31^. However, these interactions have not previously been studied in all possible combinations of steroid hormone receptor crosstalk, leaving interactions of potential clinical consequence unexplored. Here, we developed a panel of optimized steroid hormone receptor-expressing plasmids for use in NanoBRET assays to rapidly quantify receptor homo- and heterodimerization in a live-cell, scalable setting.

Using the NanoBRET platform, we identified a PR ligand-responsive, proximity-based interaction between ERα and PR, potentially indicative of heterodimer formation (Fig. 5b). Given previous research investigating the role of ERα/PR crosstalk in breast cancer^12, 17, 31^, we created an ERα NanoBRET plasmid harboring the ET resistance-associated ERα Y537S mutation to determine if ERα/PR proximity-based interaction is altered in the context of ERα Y537S. ERα Y537S and PR proximity-based interaction was significantly induced by PR stimulation with R5020, and to a much greater extent than with ERα WT and PR (Fig. 5c). These findings not only support the value of the NanoBRET method for investigating nuclear receptor heterodimerization but also indicated a potential PR-driven ERα/PR heterodimerization that is enhanced by the ERα Y537S mutation.

## Conclusion

In summary, we developed a valuable panel of nuclear receptor expression plasmids specifically for use in NanoBRET assays to assess nuclear receptor proximity, including that of ERα and PR through ERα/PR crosstalk. We identified increased proximity-based interaction between ERα and PR in the context of the constitutively activating ERα Y537S mutation. These findings suggest a role of the ERα Y537S mutation beyond that of constitutive activity of the receptor; it also impacts ERα/PR crosstalk and thus may indicate dependence on alternative signaling pathways. It is anticipated that future research to identify such pathways altered by ERα/PR crosstalk in the context of the ERα Y537S mutation will likely identify potential targetable sensitivities in treatment-resistant ERα-positive breast cancer.

## List of Abbreviations

AF-2: activating function domain 2
AI: Aromatase inhibitor
Aldo: Aldosterone
AR: androgen receptor
ATCC: American Type Culture Collection
CSS: charcoal-stripped fetal bovine serum
DBD: DNA binding domain
DHT: 5α-dihydrotestosterone
DMEM: Dulbecco’s Modified Eagle Medium
DNA: deoxyribonucleic acid
E2: β-estradiol
ERα: estrogen receptor alpha
*ESR1*: estrogen receptor gene
ET: endocrine therapy
FBS: fetal bovine serum
LBD: ligand binding domain
MR: mineralocorticoid receptor
P4: progesterone
PBS: phosphate buffered saline
PCR: polymerase chain reaction
Pen/Strep: penicillin and streptomycin
*PGR*: progesterone receptor gene
PPAR-ϒ: peroxisome proliferator-activated receptor-ϒ
PR: progesterone receptor
PR-B: progesterone receptor, B isoform
RXR: retinoid X receptor
SERDs: selective estrogen receptor degraders
SERMs: selective estrogen receptor modulators

## Availability of data and materials

The datasets used during the current study are available from the corresponding author upon reasonable request.

## Authors’ contributions

RJH, DH, and GLG contributed to the conception and design of the work presented in this publication. Data were acquired and analyzed by RJH, DH, AIO, KL, and ETT. Data was interpreted by RJH, DH, and AIO. RJH drafted the work, with substantial revisions by RJH, DH, AIO, and GLG. RJH, DH, AIO, KL, ETT, and GLG approve the submitted version and agree to be personally accountable for the author’s own contributions and to ensure that questions related to the accuracy or integrity of any part of the work are appropriately investigated, resolved, and the resolution documented in the literature. All authors read and approved the final manuscript.

## Acknowledgments

NanoBRET assays were completed at the University of Chicago Cellular Screening Center (CSC, RRID: SCR_017914).

## Supplemental Materials

**Supplemental Table 1.**
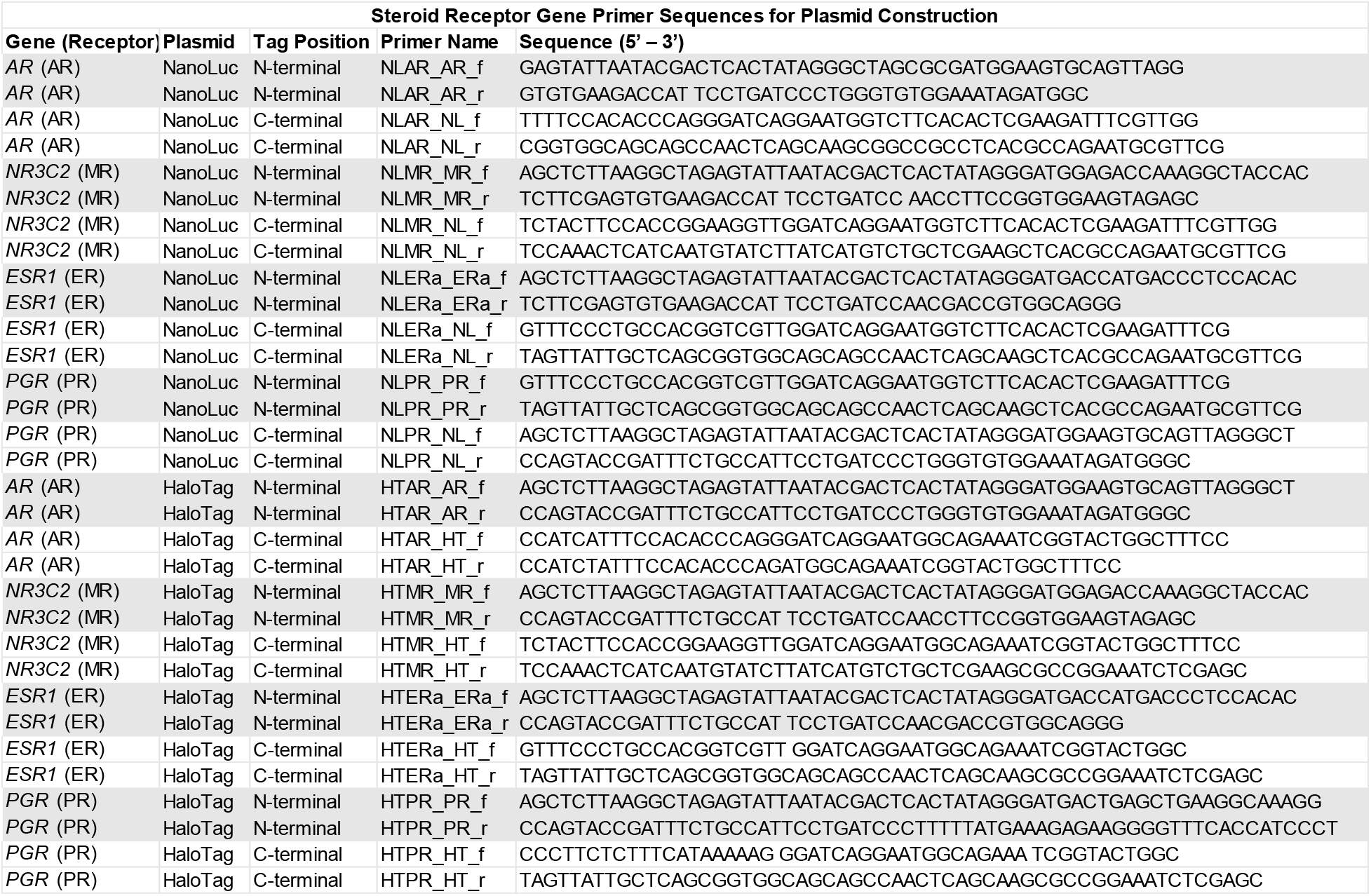
Steroid Receptor Gene Primer Sequences for Plasmid Construction. Primer sequences used for designing hormone receptor expression plasmids for NanoBRET assay.

**Supplemental Figure 1.**
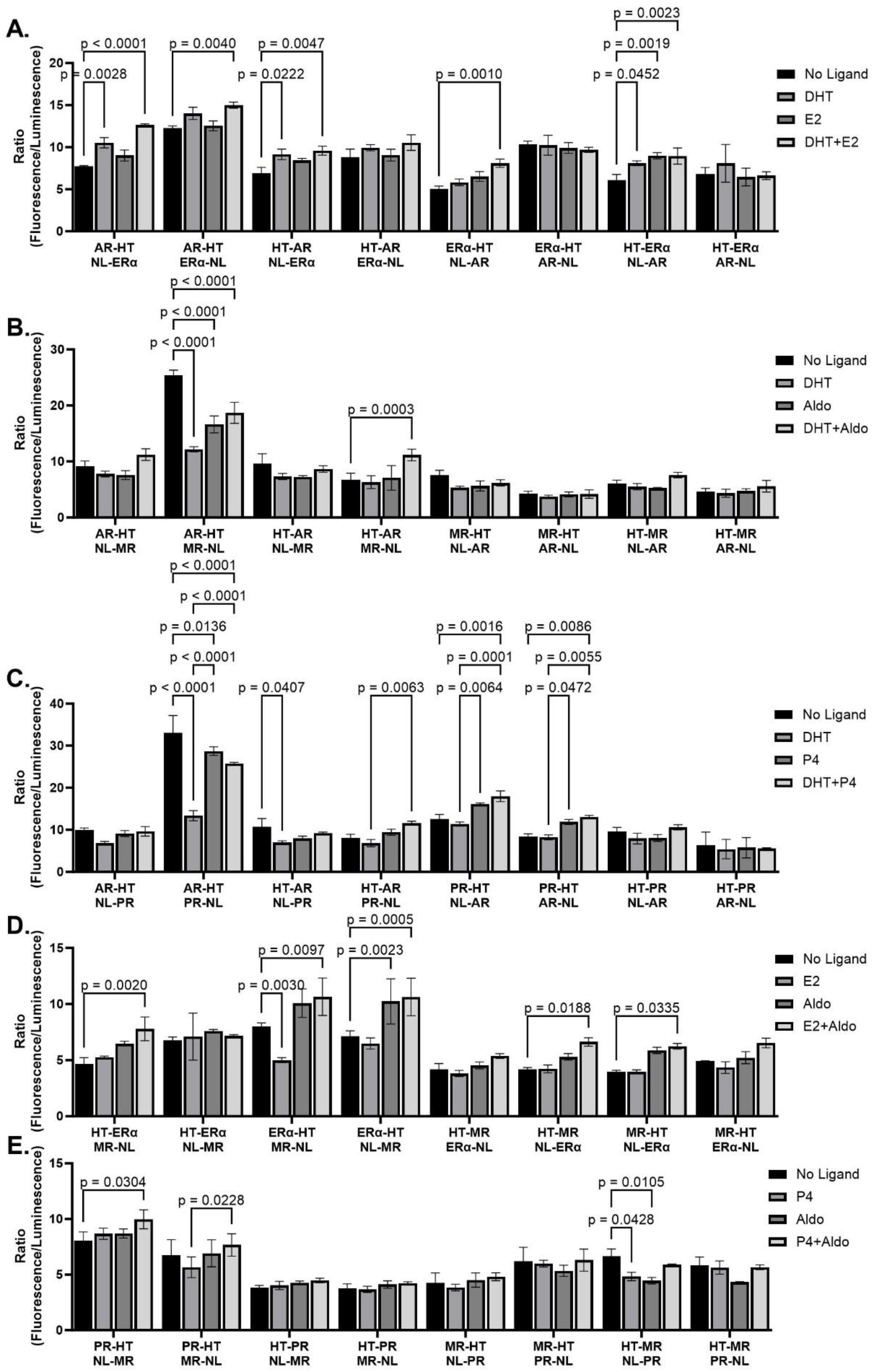
Optimization of HaloTag and NanoLuc position relative to nuclear receptor conformation for heterodimerization. A-E. NanoBRET ratios of fluorescence to luminescence for each combination of HaloTag and NanoLuc conformations to measure proximity-based interaction of **A)** AR and ERα in response to vehicle, DHT (AR native ligand), E2 (ERα native ligand), or both, **B)** AR and MR in response to vehicle, DHT, Aldo (MR native ligand), or both, **C)** AR and PR in response to vehicle, DHT, P4 (PR native ligand), or both, **D)** ERα and MR in response to E2, Aldo, or both, and **E)** MR and PR in response to vehicle, P4, Aldo, or both. Data from 3 replicates.

